# Specific targeting of intestinal *Prevotella copri* by a *Listeria monocytogenes* bacteriocin

**DOI:** 10.1101/680801

**Authors:** Nathalie Rolhion, Benoit Chassaing, Marie-Anne Nahori, Jana de Bodt, Alexandra Moura, Marc Lecuit, Olivier Dussurget, Marion Bérard, Massimo Marzorati, Hannah Fehlner-Peach, Dan R. Littman, Andrew T. Gewirtz, Tom Van de Wiele, Pascale Cossart

## Abstract

Deciphering the specific function of every microorganism in microbial gut communities is a key issue to interrogate their role during infection. Here, we report the discovery of a *Listeria* bacteriocin, Lmo2776, that specifically targets the abundant gut commensal *Prevotella copri* and affects *Listeria* infection. Oral infection of conventional mice with a Δ*lmo2776* mutant leads to a thinner intestinal mucus layer and higher *Listeria* loads both in the intestinal content and deeper tissues compared to WT *Listeria*, while no difference is observed in germ-free mice. This microbiota-dependent effect is phenocopied by precolonization of germ-free mice before *Listeria* infection, with *P. copri*, but not with other commensals,. Together, these data unveil a role for *Prevotella* in controlling intestinal infection, highlighting that pathogens may selectively deplete microbiota to avoid excessive inflammation.

## Introduction

*Prevotella* is classically considered a common commensal bacterium due to its presence in several locations of the healthy human body, including the oral cavity, gastrointestinal tract, urogenital tract and skin (*1*). The *Prevotella* genus encompasses more than 40 different culturable species of which three, *P. copri, P salivae* and *P. stercorea*, can be isolated from the gut. *Prevotella* has been reported to be associated with opportunistic infections, e.g. periodontitis or bacterial vaginosis (*1*). Moreover, *Prevotella* is the major genus of one of the three reported human enterotypes (*2*), but how *Prevotella* behaves in different gut ecosystems and how it interacts with other bacteria of the microbiota and/or with its host is not well defined. In addition, high levels of genomic diversity within *Prevotella* strains of the same species have been observed (*3*), which adds another layer of complexity for predicting the effects of *Prevotella* strains. Recent studies have linked higher intestinal abundance of *P. copri* to rheumatoid arthritis (*4-6*), metabolic syndrome (*7*), low-grade systemic inflammation (*7*) and inflammation in the context of human immunodeficiency virus (HIV) infection (*8-10*), suggesting that some *Prevotella* strains may trigger and/or worsen inflammatory diseases (*1, 11, 12*)

The microbiota plays a central role in protecting the host from pathogens, in part through a process referred to as colonization resistance (*13*). In the case of *Listeria monocytogenes*, the foodborne pathogen responsible for listeriosis, the intestinal microbiota provides protection, as germfree mice are more susceptible to infection than conventional mice (*14, 15*). Treatment with probiotics such as *Lactobacillus paracasei* CNCM I-3689 or *Lactobacillus casei* BL23 was shown to decrease *L. monocytogenes* systemic dissemination in orally inoculated mice (*16*). Unravelling the interactions between the host, the microbiota and pathogenic bacteria is critical for the design of new therapeutic strategies *via* manipulation of the microbiota. However, identifying the specific molecules and mechanisms used by the commensals to elicit their beneficial action is challenging due to the high complexity of the microbiome, together with technical issues in culturing many commensal species. In addition, cooperative interactions between commensal species are likely to be central to the functioning of the gut microbiota (*17*). So far, the mechanism or the molecules underlying the impact of commensals on the host have been elucidated only for a few species. Segmented filamentous bacteria (SFB) were shown to coordinate maturation of T cell responses towards Th17 cell induction (*18, 19*). Glycosphingolipids produced by the common intestinal symbiont *Bacteroides fragilis* have been found to regulate homeostasis of host intestinal natural killer T cells (*20*). A polysaccharide A (PSA) also produced by *B. fragilis* induces and expands Il-10 producing CD4+ T cells (*21-23*). Finally, the microbial anti-inflammatory molecule (MAM) secreted by *Faecalibacterium prausnitzii* impairs the nuclear-factor (NF)-𝒦B pathway (*24*).

Conversely, enteric pathogens have evolved various means to outcompete other species in the intestine and access nutritional and spatial niches, leading to successful infection and transmission (*25, 26*). In this regard, the contribution of bacteriocins and type VI secretion system effectors during pathogen colonization of the gut is an emerging field of investigation. Here, by studying the impact of a novel *L. monocytogenes* bacteriocin (Lmo2776) on infection, we discovered *P. copri*, an abundant gut commensal, as the primary target of Lmo2776 in both the mouse and human microbiota and as a modulator of infection.

## Results

### Lmo2776 limits *Listeria* intestinal colonization and virulence in a microbiota-dependent manner

A recent reannotation of the genome of the *Listeria monocytogenes* strain EGD-e revealed that the *lmo2776* gene, absent in the non-pathogenic *Listeria innocua* species (**Figure S1A**), potentially encodes a secreted bacteriocin of 107 amino acids (*27, 28*), homologous to the lactococcin 972 (Lcn972) secreted by *Lactococcus lactis* (*29*) and to putative bacteriocins of pathogenic bacteria *Streptococcus iniae* (*30*), *Streptococcus pneumoniae* and *Staphylococcus aureus* (**Figure S1B**). This gene belongs to a locus containing two other genes *lmo2774* and *lmo2775* genes, encoding potential immunity and transport systems (*28*). This locus is present in Lineage I strains responsible for the majority of *Listeria* clinical cases (*31*) and in some Lineage II strains, such as EGD-e (**Figure S1C**). Little is known about this bacteriocin family and most studies have focused on Lcn972. Lmo2776 shares between 38 to 47% overall amino acid sequence similarity with members of the lactococcin 972 family. Because expression of *lmo2774, lmo2775* and *lmo2776* genes is significantly higher in stationary phase compared to exponential phase of EGD-e at 37 °C in BHI (**Figure S2A**), all experiments described below were conducted with *Listeria* grown up to stationary phase.

We first examined the effect of Lmo2776 on infection. We inoculated conventional BALB/c mice with either the WT, the Δ*lmo2776* or the Lmo2776 complemented strains and compared *Listeria* loads in the intestinal lumen and deeper organs, the spleen and liver. We had verified that the deletion of *lmo2776* was not affecting the expression of surrounding genes, *lmo2774, lmo2775* and *lmo2777* (**Figure S2B**) or bacterial growth *in vitro* (**Figure S2C**). Inoculation of mice with Δ*lmo2776* strain resulted in significantly higher bacterial loads in the small intestinal lumen 24h post-inoculation compared to the WT strain (**Figure 1A**). These differences persisted at 48 and 72h post-inoculation (**Figure S2D**). Bacterial loads of Δ*lmo2776* were also significantly higher in the spleen and liver at 72h post-inoculation compared with both WT and Lmo2776-complemented strains (**Figure 1B and C**). Similar results were observed in C57BL/6J mice (data not shown). Together, these results indicate a key role for Lmo2776 in bacterial colonization of the intestine and deeper organs. Following intravenous inoculation of BALB/c mice with 5.10^3^ WT or Δ*lmo2776* bacteria, bacterial loads at 72h post-inoculation were similar in the spleen and liver (**Figure S2E**), revealing that Lmo2776 exerts its primary role during the intestinal phase of infection and not later.

**Figure 1.**
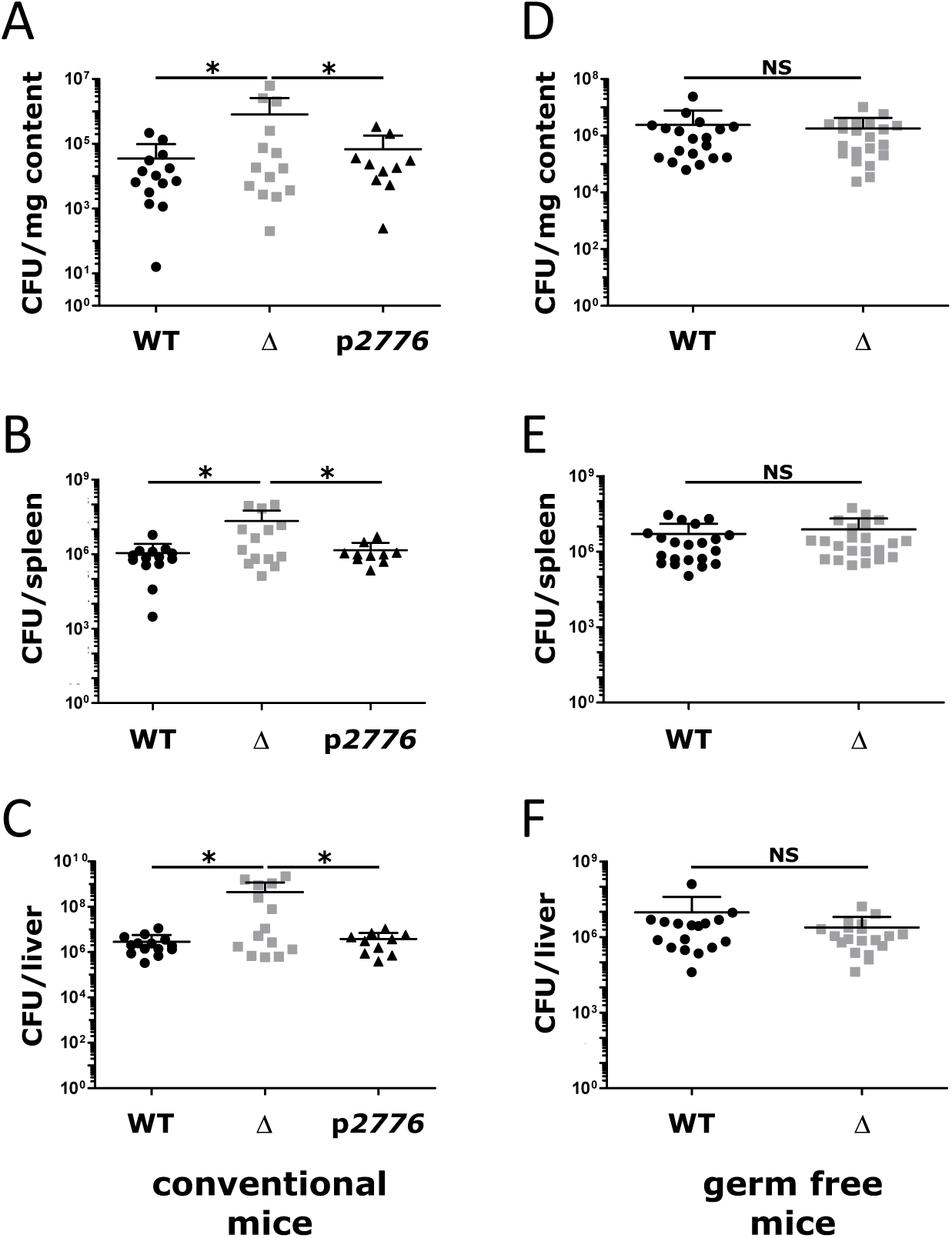
Lmo2776 limits *Listeria* virulence in a microbiota-dependent manner. (**A-C**) BALB/c mice were inoculated orally with 5×10^9^ *Listeria monocytogenes* WT (EGDe), Δ*lmo2776* or Lmo2776 complemented (*p2776*) bacteria. CFUs in the intestinal luminal content (**A**), the spleen (**B**) and the liver (**C**) were assessed at 72h post-infection. (**D-F**) Germ-free C57BL/6J were inoculated with 5×10^9^ *Listeria* WT or Δ*lmo2776* for 72h and CFUs in the intestinal luminal content (**D**), the spleen (**E**) and the liver (**F**) were assessed. Each dot represents the value for one mouse. Statistically significant differences were evaluated by the Mann–Whitney test. (*p< 0.05, NS, not significant).

Considering that *lmo2776* is predicted to encode a bacteriocin and that it significantly affects the intestinal phase of infection, we hypothesized that Lmo2776 might target intestinal bacteria, thereby impacting *Listeria* infection. To address the role of intestinal microbiota in infection, we orally inoculated germ-free mice with WT or Δ*lmo2776* strains and compared bacterial counts 72h post-inoculation. Strikingly, no significant difference was observed between WT and Δ*lmo2776* strains in the small intestinal content (**Figure 1D**), nor in spleen and liver (**Figure 1E and F**). These results showed that the Lmo2776 bacteriocin limits the virulence of wild-type *Listeria* in a microbiota-dependent manner.

### Lmo2776 specifically targets *Prevotella* in mouse and human microbiota

In order to identify which intestinal bacteria were targeted by Lmo2776, we compared microbiota compositions of conventional mice orally infected with WT or Δ*lmo2776* strains by 16S rRNA gene sequencing. We first verified that the fecal microbiota composition of all mice was indistinguishable at day 0 (**Figure 2A**). As expected, the microbiota composition at day 1 post-infection was dramatically altered by infection with *Listeria* WT (**Figure S3A**). These alterations in microbiota composition included reduced levels of *Bacteroidetes* phylum (relative abundance before infection: 65.4% and at day 1 post-infection: 42.4%) and increased levels of *Firmicutes* (relative abundance before infection: 29.9% and at day 1 post infection: 54.0%) (**Figure S3B to E**). The increased levels in the *Firmicutes* were mainly due to an increase of the *Clostridia* class (relative abundance before infection: 27.4% and at day 1 post-infection: 50.7%). Of note, the relative abundance of *Listeria* was around 0.1% and cannot therefore explain by itself the increased levels of *Firmicutes* observed between day 0 and day 1. Importantly, at 24h and 48h post-infection, intestinal microbial community compositions differed in mice orally inoculated with the Δ*lmo2776* strain compared to the WT strain (**Figure 2A**). We focused on operational taxonomic units (OTUs) for which the relative abundance was identical before the infection with the *Listeria* strains (day -3 to day 0) and was subsequently altered by at least a 2-fold difference at day 1 post-infection in mice infected with Δ*lmo2776* compared to mice infected with WT strain. In independent experiments, the relative abundance of 12 OTUs was lower in mice infected with the WT strain compared to the Δ*lmo2776* mutant (**Figure 2B and C**) (OTU 355746, 216524, 421792, 258849, 331772, 346870, 430194, 447141, 465433, 208409, 353012 and 364179) at day 1 and also at day 2 post-infection. Phylogenetic analyses revealed that all these 12 OTUs belong to the *Prevotella* cluster (**Figure 2D**). A decrease of *Prevotella* in mice infected with WT strain at day 1 and day 2 post-inoculation compared to mice infected with Δ*lmo2776* strain was also observed by qPCR analysis, using primers specific for *Prevotella*, confirming that Lmo2776 targets *Prevotella* in the intestinal microbiota (**Figure 2E**).

**Figure 2.**
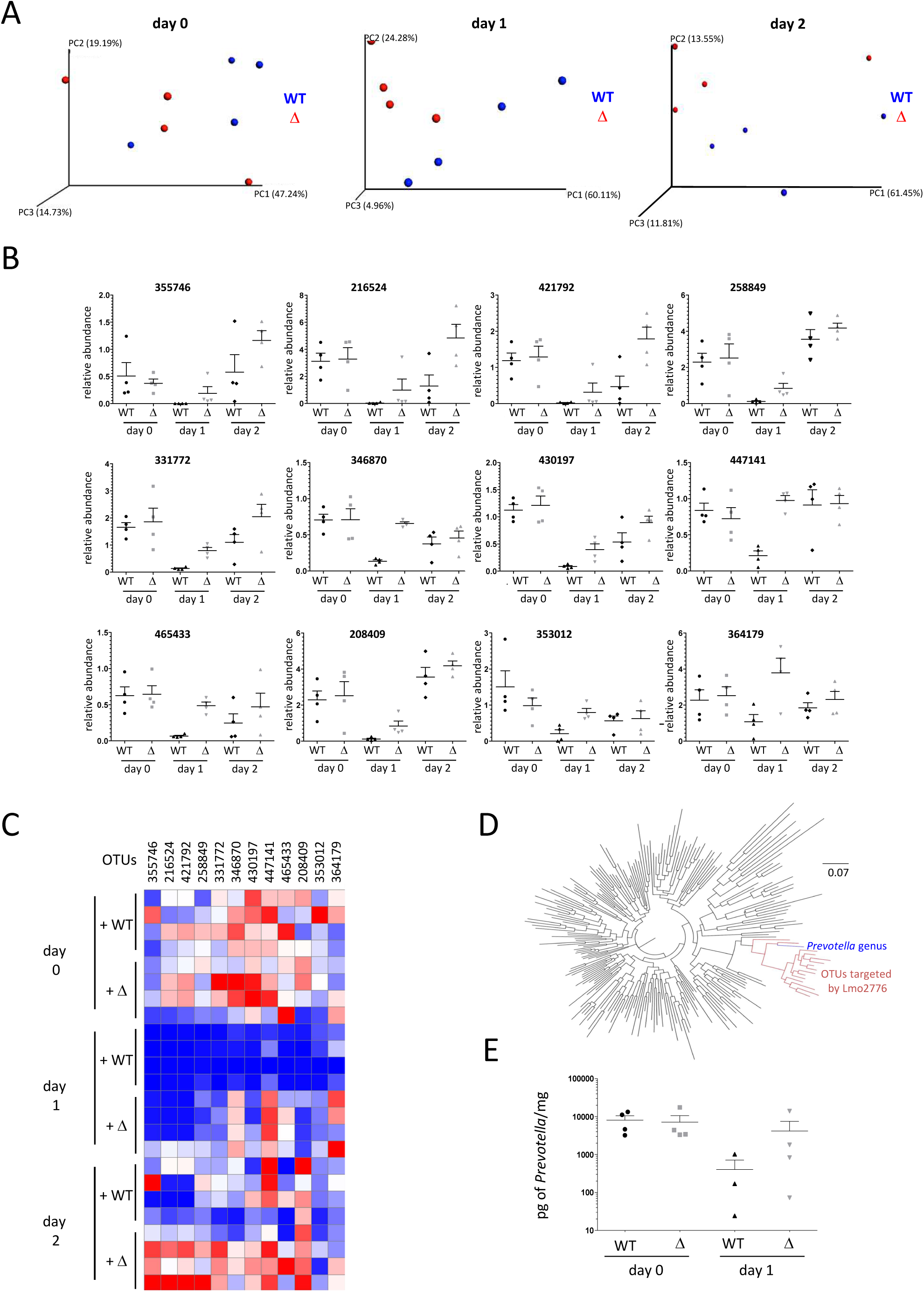
Lmo2776 targets *Prevotella* in mouse microbiota. (**A**) Principal coordinates analysis of the weighted Unifrac distance matrix of mice infected with WT strain (blue) or Δ*lmo2776* (red) at day 0 (left), day 1 (center) and day 2 (right). Permanova: at day 0, P=0.383; at day 1, P=0.05864; and at day 2, P=0.360. (**B**) Relative abundance of 12 OTUs in gut microbiota of mice inoculated with WT or Δ*lmo2776* strains at day 0, day 1 and day 2. Each dot represents the value for one mouse. (**C**) Heat-map analysis of the relative abundance of 12 OTUs in gut microbiota of mice inoculated with WT or Δ*lmo2776* strains at day 0, day 1 and day 2. Each raw represents one mouse. The red and blue shades indicate high and low abundance. (**D**) Phylogenetic tree of 16S rRNA gene alignment of 5 representative bacteria for each phylum of the bacteria domain, together with OTUs showing significantly different relative abundances in gut microbiota of mice infected with WT or Δ*lmo2776* strains at day 1. The 12 OTUs with an increased abundance in Δ*lmo2776*-infected mice compare to *Listeria* WT-infected mice are shown in red and *Prevotella* genus is indicated in blue. (**E**) PCR quantification of *Prevotella* in feces of mice inoculated with WT strain or Δ*lmo2776* at day 0 and day 1.

Important differences exist between mouse and human gut microbiota composition. Indeed, *Prevotella* abundance is known to be low in the mouse intestinal content (less than 1%) while it can reach up to 80 % in the human gut microbiota (*32, 33*). As *Listeria* is a human pathogen, we searched to investigate the impact of Lmo2776 on human intestinal microbiota. For this purpose, we used a dynamic *in vitro* gut model (mucosal-simulator of human intestinal microbial ecosystem (M-SHIME^®^)), which allows stable maintenance of human microbiota *in vitro*, in the absence of host cells but in presence of mucin-covered beads (*34-37*) and therefore studies on human microbiota independently of the host responses (such as inflammation). The microbiota of a healthy human volunteer was inoculated to the system which was then infected with WT or Δ*lmo2776 Listeria*. Application of 16S sequencing to luminal and mucosal M-SHIME^®^ samples indicated that before *Listeria* inoculation, the bacterial composition in all vessels was similar (**Figure 3A** and data not shown). In contrast, following *Listeria* addition, luminal microbial community compositions were different in vessels containing WT bacteria compared to both non-infected vessels and vessels infected with the Δ*lmo2776* isogenic mutant (**Figure 3A**). No difference was observed in mucosal microbial community composition. The relative abundance of 7 OTUs (313121, 518820, 346938, 588929, New.0.ReferenceOTU20, 173565 and 89083) was lower in the case of the WT strain compared to the non-inoculated condition or upon addition of the Δ*lmo2776* strain (**Figure 3B**). These 7 OTUs all belonged to *Prevotella copri* species (**Figure 3C**), revealing that Lmo2776 targets *P. copri* in the human microbiota in a host-independent manner.

**Figure 3.**
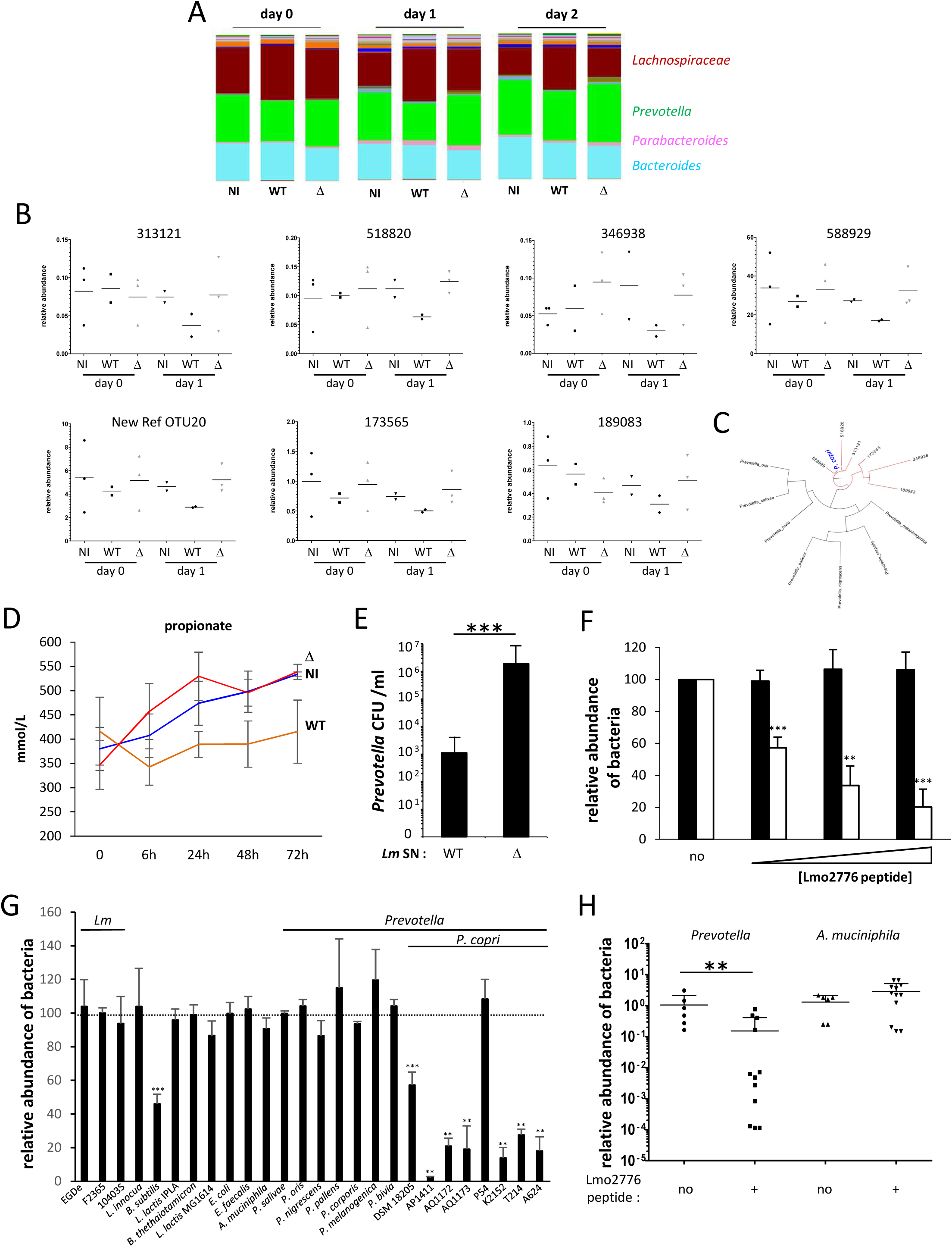
Lmo2776 targets *P. copri* in human microbiota and *in vitro*. (**A**) Relative abundance of genera in SHIME^®^ vessels non-infected or infected with WT or Δ*lmo2776* strains at day 0, day 1 and day 2. The four more abundant genera are indicated. (**B**) Relative abundance of 7 different OTUs in SHIME^®^ vessels infected with WT or Δ*lmo2776* strains or non-infected at day 0 and day 1. Each dot represents the value for one vessel. (**C**) Phylogenetic tree of 16S rRNA gene alignment of several *Prevotella* species, together with OTUs showing significantly different relative abundances in vessels inoculated with WT or Δ*lmo2776* strains at day 1. Six OTUs with an increased abundance in Δ*lmo2776*-inoculated vessels compare to *Listeria* WT-inoculated vessels are shown in red and *Prevotella copri* is indicated in blue. (**D**) Levels of propionate in SHIME^®^ vessels infected with WT (orange) or Δ*lmo2776* (red) strains or non-infected (blue) overtime. Results are expressed as mean ± SEM for 2 to 3 individual vessels. **(E)** Numbers of *P. copri* after incubation with supernatant of WT (*Lm*) or Δ*lmo2776* strains. **(F)** Relative abundance of *P. copri* (white) and *B. thetaiotamicron* (black) after 24h incubation with increasing dose of Lmo2776 peptide (3 (+), 6 (++) and 9(+++) µg/ml) relative to their abundance without Lmo2776 peptide. (**G)** Relative abundance of different bacteria after 24h incubation with Lmo2776 peptide (3µg/ml) relative to their abundance without Lmo2776 peptide. Results are expressed as mean ± SEM of a least 3 independent experiments and P-values were obtained using two-tailed unpaired Student’s t-test (*p<0.05, ***p<0.005). (**H**) PCR quantification of *Prevotella* and *A. muciniphila* in the feces of mice treated with Lmo2776 peptide (1 mg) or with water relative to their levels before treatment. Each dot represents the value for one mouse. Statistically significant differences were evaluated by Student’s t-test (**p< 0.01).

As short-chain fatty acid (SCFA) levels serve as a classical read-out for gut microbiota metabolism and as *Prevotellae* are known to produce propionate (*38*), we quantified SCFAs production in the luminal M-SHIME^®^ samples. A specific decrease in propionate production upon infection with WT bacteria was observed as early as 6h post-infection (**Figure 3D**) compared to non-infected and Δ*lmo2776*-infected vessels. This difference was continuously observed up to 3 days post-infection, while no significant difference was observed for butyrate, isobutyrate, acetate and isovalerate (**Figure S4**). Although propionate is produced by many bacterial species, the decrease in propionate production observed upon inoculation of M-SHIME^®^ with WT *Listeria* is in agreement with the decrease in *Prevotella* population.

### Lmo2776 targets *P. copri in vitro*

We first addressed the direct inhibitory activity of Lmo2776 on *P. copri* by growing *P. copri* at 37 °C in anaerobic conditions in the presence of culture supernatants of *Listeria* strains and counting the viable CFUs on agar plates. Growth of *P. copri* dramatically decreased (up to 3 Log) in the presence of the WT *Listeria* supernatant compared to the Δ*lmo2776* supernatant or medium alone (**Figure 3E**), suggesting that Lmo2776 is secreted and targets directly *P. copri*. To definitively assess the function of Lmo2776, a peptide of 63 aa (Gly69 to Lys131) corresponding to the putative mature form of Lmo2776 was synthesized. Its activity was first analyzed on *P. copri* and *B. thetaiotaomicron*, another prominent commensal bacterium (**Figure 3F**). A dose-dependent effect of Lmo2776 peptide was observed on the growth of *P. copri* while no effect was observed on the growth of *B. thetaiotaomicron*, demonstrating that Lmo2776 targets *P. copri* and not *B. thetaiotaomicron*. We then tested the effect of the peptide on several other intestinal bacteria, either aerobic (*Enterococcus faecalis, Escherichia coli*) or anaerobic (*Akkermansia muciniphila*) bacteria. No effect was observed on any of these bacteria (**Figure 3G**). Moreover, Lmo2776 peptide did not inhibit the growth of seven other *Prevotella* species (*P. salivae, P. oris, P. nigrescens, P. pallens, P. corporis, P. melaninogenica* and *P. bivia*). We next tested the peptide activity on 7 *P. copri* isolated from healthy humans and patients. Strikingly, 6 out of the 7 strains were sensitive to the bacteriocin (**Figure 3G**).

We also tested the effect of the Lmo2776 peptide on known targets of the bacteriocins of the lactococcin-972 family (*B. subtilis, L. lactis* MG1614). Growth of *B. subtilis* decreased significantly in presence of the peptide (**Figure 3G**), while no effect was observed on *L. lactis* MG1614. Growth of *Bacillus subtilis* was also specifically and significantly reduced in the presence of WT *Listeria* and of Lmo2776 complemented strains compared to the Δ*lmo2776* strain (**Figure S5A-B**). Addition of the culture supernatant of WT *Listeria* to *B. subtilis* significantly decreased the number of *B. subtilis* compared to the addition of Δ*lmo2776* culture supernatant (**Figure S5C**). (*30*). These results indicate that Lmo2776 is a *bona fide* bacteriocin that targets both *P. copri* and *B. subtilis in vitro*.

To evaluate the effect of the Lmo2776 peptide *in vivo* in animals, we used an approach previously described to bypass degradation by enzymes of the upper digestive tract. Conventional BALB/c mice were inoculated intra-rectally with Lmo2776 peptide or water, taken as a control. Levels of total bacteria, *Prevotella* and *Akkermansia muciniphila* were determined by quantitative PCR on feces collected between 1 and 4h post-administration. While no effect was observed on the levels of total bacteria or *A. muciniphila*, fecal levels of *P. copri* decreased following administration of Lmo2776 peptide, demonstrating that similar to bacteria, Lmo2776 alone was effective in reducing *P. copri in vivo* (**Figure 3H**).

### Colonization of germ-free mice by *P. copri* phenocopies the effect of the microbiota on *Listeria* intestinal growth in conventional mice

To decipher the role of *P. copri* during *Listeria* infection *in vivo*, germ-free C57BL/6J mice were orally inoculated with either *P. copri, B. thetaiotaomicron* or *P. salivae*, another *Prevotella* present in the gut, or stably colonized with a consortium of 12 bacterial species (termed Oligo-Mouse-Microbiota (Oligo-MM^12^), representing members of the major bacterial phyla in the murine gut: *Bacteroidetes* (*Bacteroides caecimuris* and *Muribaculum intestinale*), *Proteobacteria* (*Turicimonas muris*), *Verrucomicrobia* (*Akkermansia muciniphila*), *Actinobacteria* (*Bifidobacterium longum subsp. Animalis*) and *Firmicutes* (*Enterococcus faecalis, Lactobacillus reuteri, Blautia coccoides, Flavonifractor plautii, Clostridium clostridioforme, Acutalibacter muris* and *Clostridium innocuum*) (*39*)). Two weeks after colonisation, these mice were orally inoculated with WT *Listeria* or Δ*lmo2776* strains and *Listeria* loads in the intestinal lumen and target organs were compared 72h post-infection. Compared to the WT strain, the Δ*lmo2776* mutant strain displayed significantly higher loads in the intestinal lumen (**Figure 4A**), the spleen (**Figure 4B**) and liver (**Figure S5D**) in mice colonized with *P. copri*, while no difference between the two strains was observed in mice precolonized with *B. thetaiotaomicron, P. salivae* or the OligoMM^12^ consortium. In addition, the number of *P. copri* significantly decreased in WT inoculated *P. copri*-colonized animals compared to Δ*lmo2776*-inoculated animals (**Figure 4C**). Altogether, these results indicate that the greater ability of the Δ*lmo2776* mutant to grow in the intestine and reach deeper tissues compared to the WT strain is dependent on the presence of *Prevotella* in the microbiota, as it is observed in either conventional mice or mice colonized with *P. copri*.

**Figure 4.**
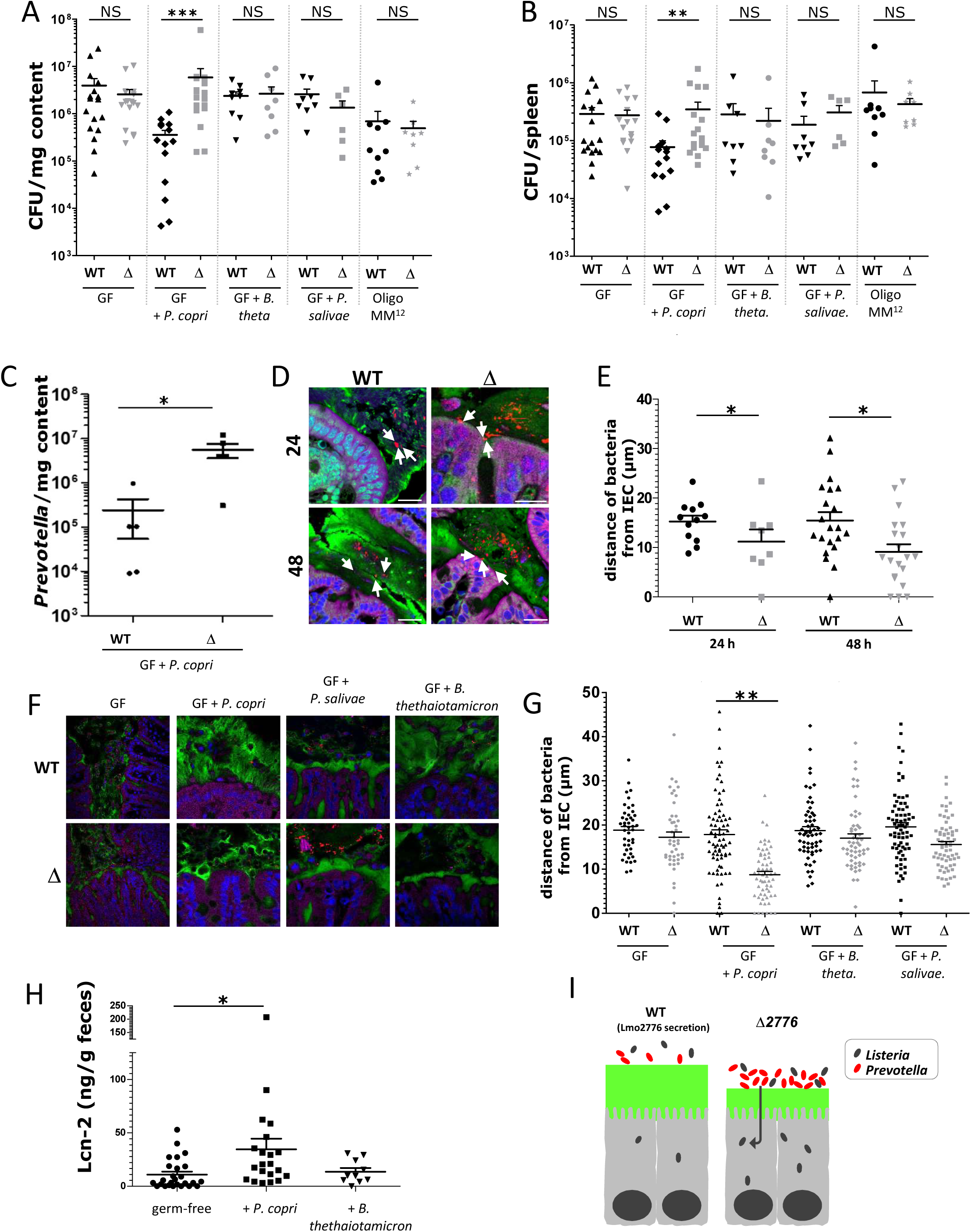
*P. copri* controls *Listeria* infection by modifying mucus layer and promoting inflammation. (**A-B**) Assessment of listerial CFUs in the intestinal luminal content (**A**) and in the spleen (**B**) of germ-free (GF) C57BL/6J mice colonized or not with *P. copri, P. salivae* or *B. thetaiotamicron* or stably colonized with 12 bacterial species (Oligo-MM^12^) for 2 weeks and then inoculated with *L. monocytogenes* WT or Δ*lmo2776* for 72h. (**C**) Numbers of *P. copri* CFUs in the intestinal luminal content of GF C57BL/6J mice colonized with *P. copri* and then inoculated with *Listeria* WT or Δ*lmo2776* for 72h. (**D**) Confocal microscopy analysis of microbiota localization in colon of BALB/c mice infected with 5×10^9^ *Listeria* WT or Δ*lmo2776* bacteria for 24 or 48h. Muc2 (green), actin (purple), bacteria (red), and DNA (Blue). Bar = 20μm. White arrows highlight the 3 closest bacteria. Pictures are representatives of 5 biological replicates. (**E**) Distances of closest bacteria to intestinal epithelial cells per condition over 5 high-powered fields per mouse, with each dot representing a measurement. **(F)** Confocal microscopy analysis of microbiota localization in colon of GF C57BL/6J mice colonized with *P. copri, B. thetaiotamicron* or *P. salivae* for 2 weeks and inoculated with 5×10^9^ *Listeria* WT or Δ*lmo2776* bacteria. Muc2 (green), actin (purple), bacteria (red), and DNA (Blue). Bar = 20μm. (**G**) Distances of closest bacteria to intestinal epithelial cells per condition over 5 high-powered fields per mouse, with each dot representing a measurement. (**H**) Levels of the inflammatory marker Lcn-2 in faeces of mice 2 weeks post-inoculation with *P. copri* or *B. thetaiotamicron.* (**I**) Model depicting the effect of *Prevotella* on *Listeria* infection. In A, B and E, each dot represents one mouse. Statistically significant differences were evaluated by the Mann–Whitney test (A, B), one way-ANOVA test (E, G) or two-tailed unpaired Student’s t-test (C and H) (*p< 0.05, ***p<0.005).

### *P. copri* modifies the mucus layer and its permeability

The intestinal mucus layer of conventional animals forms a physical barrier of about 30µm that is able to keep bacteria at a distance from the epithelium (*40*). A mucus-eroding microbiota promotes greater epithelial access (*41*). *Prevotella*, through production of sulfatases that induce mucus degradation (*42*), might impair the mucosal barrier function and therefore contribute to better accessibility to intestinal epithelial cells and to local inflammation. We thus compared the mucus layer thickness of conventional mice infected with WT *Listeria* or Δ*lmo2776* by confocal microscopy, using mucus-preserving Carnoy fixation and FISH (*43*). The average distance of bacteria from colonic epithelial cells was significantly smaller in mice infected with Δ*lmo2776* compared to mice infected with WT *Listeria* at 24 and 48h (**Figure 4D and E**), suggesting that *Prevotella* present in the microbiota of mice infected with Δ*lmo2776* decreases the mucus layer thickness and consequently increases its permeability. Of note, these distances were also smaller than in uninfected mice, indicating that *Listeria* infection by itself can affect the mucus layer thickness. To confirm the effect of *P. copri* on mucus layer in the context of listerial infection, germ-free C57BL/6J mice were precolonized with *P. copri, B. thetaiotaomicron* or *P. salivae*, then orally inoculated with WT *Listeria* or Δ*lmo2776* strains and mucus layer thickness was analysed by FISH. In mice precolonized with *P. copri*, the average distance of bacteria from colonic epithelial cells was significantly smaller in Δ*lmo2776-* infected mice compared to WT *Listeria*-infected mice (**Figure 4F and G)**. Strikingly, such difference was not observed in germ-free mice or in mice precolonized with *P. salivae* or *B. thetaiotaomicron*, revealing that mucus erosion is dependent on *P. copri*.

Since disruption of the mucosal barrier by *Prevotella* could favour invasion of the host by bacteria and contribute to intestinal inflammation, we quantified faecal lipocalin-2 (LCN-2) as a marker of intestinal inflammation (*44*). LCN-2 is a small secreted innate immune protein which is critical for iron homeostasis and whose levels increase during inflammation. Faecal LCN-2 levels were thus analysed after colonization of germ-free mice with *P. copri* compared to non-colonized mice or mice colonized with *B. thetaiotaomicron*. A significant increase of faecal LCN-2 was observed in germ-free mice monocolonized with *P. copri* compared to non-colonized animals or to animals monocolonized with *B. thetaiotaomicron* (**Figure 4H**), revealing that *P. copri* induces intestinal inflammation. Altogether, these results showed that presence of *Prevotella* in the intestine is associated with a thinner mucus layer and increased levels of faecal LCN-2. They are consistent with previous reports describing *Prevotella* as a bacterium promoting a pro-inflammatory phenotype (*1, 6, 45*).

## Discussion

Outcompeting intestinal microbiota stands among the first challenging steps for enteropathogens. Pathogens may secrete diffusible molecules such as bacteriocins or T6SS effectors to target commensals and consequently promote colonization and virulence. In most cases, the molecular mechanisms underlying the interplay between pathogenic and commensal bacteria in the intestine remain elusive. We previously reported that most strains responsible for human infections, such as the F2365 strain, secrete a bacteriocin that promotes intestinal colonization by *Listeria* (*46*). When overexpressed in mouse gut, this bacteriocin, named Listeriolysin S (LLS), decreases *Allobaculum* and *Alloprevotella* genera known to produce butyrate or acetate, two SCFAs reported to inhibit transcription of virulence factors or growth of *Listeria* (*47, 48*). However, whether physiological concentrations of LLS have a direct or an indirect role on these *genera* is still under investigation and is a question particularly difficult to address as LLS is highly post-translationally modified and therefore difficult to purify or to synthetize. In the case of *Salmonella enterica* serovar Typhimurium infection, killing of intestinal *Klebsiella oxytoca via* the its T6SS is essential for *Salmonella enterica* gut colonization of gnotobiotic mice colonized by *K. oxytoca* (*49*), but whether *K. oxytoca* and other members of the gut microbiota are targeted by the *Salmonella* T6SS in conventional mice is unknown. Finally, *Shigella sonnei* uses a T6SS to outcompete *E. coli in vivo* but the effectors responsible for this effect are unknown (*50*).

Here, we demonstrated that the Lmo2776 *Listeria* bacteriocin targets *Prevotella* in mouse and in *in vitro* reconstituted human microbiota. This effect is direct and specific to *P. copri* as (*i*) *P. copri* are killed by *Listeria* culture supernatant and by the purified Lmo2776 *in vitro* and (*ii*) despite the complexity of the microbiota and its well-controlled equilibrium, no other genus of the intestinal microbiota was found to be impacted by Lmo2776. By studying Lmo2776, we have unveiled a so far unknown role for intestinal *Prevotella copri* in controlling bacterial infection. The intestinal microbiota, in some cases, has already been reported to promote bacterial virulence by producing metabolites that enhance pathogens virulence gene expression and colonization in the gut (*25, 26*). For example, *B. thetaiotaomicron* enhances *Clostridium rodentium* colonization by producing succinate (*51, 52*) and *Akkermansia muciniphila* exacerbates *S. Typhimurium*-induced intestinal inflammation by disturbing host mucus homeostasis (*53*). *P. copri* increases the mucus layer permeability and increases propionate concentration and levels of fecal LCN-2, in agreement with previous studies reporting that *P. copri* exacerbates inflammation (*6, 45*). In addition, *Prevotella* enrichment within the lung microbiome of HIV-infected patients has been observed and is associated with Th17 inflammation (*54*). *Prevotella* sp. have also been associated with bacterial vaginosis and their role in its pathogenesis has been linked to the production of sialidase, an enzyme involved in mucin degradation and increased levels of pro-inflammatory cytokines (*55, 56*). Our data strongly indicate that *P. copri*, by modifying the mucus layer permeability and changing the gut inflammatory condition, promotes greater epithelial access and therefore infection by *Listeria* (**Figure 4I**). We can speculate that individuals with high abundance of intestinal *Prevotella* might be more susceptible to enteric infections. Interestingly, it was recently shown that subjects with higher relative abundance of *P. copri* could be at higher risk to traveler’s diarrhea and to the carriage of multidrug-resistant *Enterobacteriaceae* (*57*). On the other hand, Lmo2776 *Listeria* bacteriocin allows a selective depletion of *P. copri* in intestinal microbiota. This could therefore prevent excessive inflammation and allow *Listeria* persistence and long lasting infection, eventually leading to menngitis. Further work is required to determine why *Listeria* strains would gain an advantage by keeping the *lmo2776* gene. We showed here that the Lmo2776 bacteriocin also targets *B. subtilis*, a Gram-positive bacterium found in the soil, suggesting that Lmo2776 could give an advantage to *Listeria* in that environment. It is possible that Lmo2776 is critical for species survival and replication in a so far unknown niche, consequently favoring transmission or dissemination. *B. subtilis* is also found in the human gastrointestinal tract (*58*) and could be targeted by Lmo2776 in the intestine as well. *B. subtilis* is also targeted by Sil, another member of the lactococcin 972 family (*30*). The role of the homologs of lactococcin 972 in other human pathogenic bacteria such as *S. pneumoniae* and *S. aureus* remains to be determined, but the conservation of the bacteriocin in different pathogenic bacteria associated with mucosa strongly suggests an important role.

Taken together, our data indicate that *P. copri* can modulate human disease and using Lmo2776 might represent an effective therapeutic tool to specifically reduce *P. copri* abundance in the gut without affecting the remaining commensal microbiota.

## Acknowledgments

We thank the CDTA (Cryopréservation, Distribution, Typage et Archivage animal, CNRS, Orléans) and the Institut Pasteur Animalerie Centrale staff, especially the technicians of the Centre for Gnotobiology Platform of their help with animal work (Karim Sébastien, Thierry Angélique, Marisa Gabriela Lopez Dieguez, Martine Jacob and Eddie Maranghi). We thank Grégory Jouvion and Magali Tichit from the Unité d’Histopathologie humaine et modèles animaux de l’Institut Pasteur for technical help and the Centre Ressources Biologiques de l’Institut Pasteur for providing strains. We thank Laurence Maranghi and Juienne Blondou for essential technical support and Christophe Bécavin, Andrew Hryckowian, Beatriz Martinez, Gérard Eberl and Gunnar Hansson for helpful discussion.

N.R. was supported by an EMBO short-term fellowship (ASTF 399-2015). B.C. is supported by a Career Development Award from the Crohn’s and Colitis Foundation, an Innovator Award from the Kenneth Rainin Foundation and a Seed Grant from the GSU’s Brain & Behaviour program. O.D. was supported by Idex UP2019. H. F-P. was supported by an NYU-HCC CTSI grant (1TL1TR001447). D.R. L. was supported by the Howard Hughes Medical Institute and the Colton Center for Autoimmunity. This work was supported by grants to P.C. (European Research Council (ERC) Advanced Grant BacCellEpi (670823), ANR BACNET (BACNET 10-BINF-02-01), ANR Investissement d’Avenir Programme (10-LABX-62-IBEID), Human Frontier Science Program (HFSP; RGP001/2013), ERANET Infect-ERA PROANTILIS (ANR-13-IFEC-0004-02) and the Fondation le Roch les Mousquetaires). P.C. is a Senior International Research Scholar of the Howard Hughes Medical Insitute.

## List of supplementary materials

Materials and Methods Fig S1-S5

References(59-72)

