## supplemental material for "Specific targeting of intestinal *Prevotella copri* by a *Listeria monocytogenes* bacteriocin"

#### Supplementary materials

#### Methods

##### Bacterial strains and plasmids

Strains, plasmids and primers used in this study are listed in the tables S1 and S2 respectively. For standard experiments, *Listeria*, *E. coli*, *B. subtilis* and *L. lactis* were grown at 37°C with shaking in Brain Heart Infusion (BHI) medium (Difco) and Luria-Bertani (LB) medium (BD). If needed, *Lm* were selected on Oxford medium (Oxoid). Anaerobic bacteria were grown in appropriate medium (PYG medium modified or Schaedler medium or BHI supplemented with 8mM L-Cysteine hydrochloride, 0.2% $NaHCO_3$ and 0.025% Hemin, following ATCC or DSMZ recommendations) at 37°C under anaerobic conditions (Genbag Anaer, Biomérieux or AnaeroGen, ThermoScientific). The Lmo2776 deletion mutant was constructed using the pMAD shuttle plasmid (*59*) as described previously (*60*) and confirmed by DNA sequencing. For the construction of pAD-based plasmid, fragment obtained by PCR with EGD-e genomic DNA as template, was cloned into the SmaI/SalI sites of the pAD vector (*61*) derived previously from the pPL2 vector (*62*). Plasmid was verified by sequencing with primers pPL2-Rv and pPL2-Fw and were transformed into Δ*lmo2776* by electroporation. Integration in the chromosome was verified by PCR using primers NC16 and PL95 (*61*). *P. copri* strains (AP1411, AQ1172, AQ1173, P54, K2152, T214 and A624) were isolated from stool from healthy subjects and new onset rheumatoid arthritis patients. Stool was collected into anaerobic transport media (Anaerobe Systems), then streaked on BRU and LKV plates (Anaerobe Systems). After 24-48h, individual colonies were picked and screened with *Prevotella*-specific PCR primers, and the 16S rRNA V3-V4 sequence was confirmed by Sanger sequencing (Fehlner-Peach *et al.*, manuscript in preparation). *Prevotella*-positive isolates were grown on BRU plates, and glycerol stocks were frozen at -80°C.

##### Mice

9- to 12-week-old female BALB/c conventional (Charles River) or C57BL6/J conventional (Charles River) or C57BL6/J germfree (CDTA or Pasteur) or C57BL6/J Oligo-$MM^{12}$ (*39*) wild-type mice were used for experiments. Germfree and Oligo-$MM^{12}$ mice were housed in plastic gnotobiotic isolators.

All animal experiments were carried out in strict accordance with the French national and European laws and conformed to the Council Directive on the approximation of laws, regulations, and administrative provisions of the Member States regarding the protection of animals used for experimental and other scientific purposes (86/609/Eec). Experiments that relied on laboratory animals were performed in strict accordance with the Institut Pasteur's regulations for animal care and use protocol, which was approved by the Animal Experiment Committee of the Institut Pasteur (approval no. 03-49).

**Mice infection**

*Lm* overnight cultures were diluted in BHI and bacteria were grown to an optical density at 600 nm (OD600) of 1. Bacterial cultures were centrifuged at 3,500 × g for 15 min and washed three times in phosphate-buffered saline (PBS). Mice were infected orally with 5 × $10^9$ bacteria diluted in 200 µl of PBS supplemented with 300 µl of $CaCO_3$ (50 mg/ml). Serial dilutions of the inoculum were plated to control the number of bacteria inoculated. The different inoculum were closed to 5 x $10^9$ bacteria and the mean ± SEM of all independent experiments were : for WT : 5.03 x $10^9$ ± 0.26 x $10^9$, for Δ*lmo2776* : 5.01 x $10^9$ ± 0.14 x $10^9$ (P= 0.45) or Lmo2776 complemented strain : 4.87 x $10^9$ ± 0.30 x $10^9$ (P= 0.25). Mice were killed at subsequent time points, and intestines, spleens, and livers were removed. The small intestine was opened, and the luminal content was recovered in a 1.5-mL tube and weighed. The small intestine (duodenum, jejunum, and ileum) tissue was washed four times in DMEM (ThermoFisher), incubated for 2 h in DMEM supplemented with 40 µg/mL gentamycin, and finally washed four times in DMEM. All of the organs and the intestinal luminal content were homogenized, serially diluted, and plated onto BHI plates or Oxford plates and grown overnight at 37 °C for 48–72 h. CFU were enumerated to assess bacterial load. At least three independent experiments were carried out with four or five mice per group in each experiment. Statistically significant differences were evaluated by the Mann–Whitney test, and differences were considered statistically significant when P values were <0.05.

For mice colonisation, *P. copri*, *P. salivae* and *B. thetaiotamicron* were grown to log phase under anaerobic conditions in PYG modified or Schaedler broth media and $10^7$ CFU were used to inoculate germ-free mice. Feces were collected at 2 weeks post-gavage to confirm colonization. Feces were homogenized, serially diluted and plated on PYG modified, Schaedler or Columbia agar plates.

Six- to 8-week-old female BALB/c mice (Charles River, Inc., France) were injected intravenously with 5.$10^3$ CFU of the indicated strain. Mice were sacrificed at 72 h after infection, and livers and spleens were removed. Organs were homogenized and serially diluted. Dilutions were plated onto BHI plates and grown during 24 h at 37°C. Colonies were counted to assess bacterial load per organ.

**Fecal microbiota analysis by 16S rRNA gene sequencing using Illumina technology**

Before infection, 8 BALB/c conventional mice were co-housed for 5 weeks in order to stabilize and homogenize their microbiota. After oral infection, animals were single-housed. Feces were collected and frozen at -20°C. 16S rRNA gene amplification and sequencing were done using the Illumina MiSeq technology following the protocol of Earth Microbiome Project with their modifications to the MOBIO PowerSoil DNA Isolation Kit procedure for extracting DNA (www.earthmicrobiome.org/emp-standard-protocols) (*63, 64*). Bulk DNA were extracted from frozen extruded feces using a PowerSoil DNA Isolation kit (MoBio Laboratories) with mechanical disruption (bead-beating). The 16S rRNA genes, region V4, were PCR amplified from each sample as described in Chassaing *et al*., 2015 (*65*). Four independent PCRs were performed for each sample, combined, purified with Ampure magnetic purification beads (Agencourt), and products were visualized by gel electrophoresis. Products were then quantified (BIOTEK Fluorescence Spectrophotometer) using Quant-iT PicoGreen dsDNA assay. A master DNA pool was generated from the purified products in equimolar ratios. The pooled products were quantified using Quant-iT PicoGreen dsDNA assay and then sequenced using an Illumina MiSeq sequencer (paired-end reads, 2 × 250 bp) at Cornell University (Ithaca, USA).

**16S rRNA gene sequence analysis**

Analysis of the 16S rRNA gene sequence was performed exactly as previously described (*65*). Our full 16S rRNA gene sequence data are deposited under Study ID XXXX in the QIIME-DB database (http://www.microbio.me/qiime/).

**Bacterial quantification in feces**

Bulk DNA were extracted from frozen extruded feces using a PowerSoil DNA Isolation kit (MoBio Laboratories) with mechanical disruption (bead-beating). q-PCR reactions were prepared with SYBR Green master mix. Reaction cycling and quantification was carried out in an C1000 touch Thermal cycler (CFX384, Biorad). Genomic DNA from *Prevotella* was used to generate a standard curve to quantitate pg of *Prevotella* present per mg of total feces.

**M-SHIME**

M-SHIME system is a dynamic *in vitro* model which simulates the lumen-associated and mucus-associated human intestinal microbial ecosystem (ProDigest, Ghent University, Belgium) (*66-68*). It consists of consecutive pH-controlled, stirred, airtight, double-jacketed glass vessels kept at 37°C and under anaerobic conditions. The system was operated as described earlier (*69*) with minor modifications The set-up used here consisted of 3 stomach-small intestine vessels and 9 proximal colon vessels in parallel (3 for non-infected condition, 3 for infection with WT and 3 for infection with Δ*lmo2776* ). The colon vessels were inoculated at the start with 40 mL fresh human faecal suspension, from a healthy volunteer with high levels of *Prevotella***,** in 500 mL sterile nutritional medium. Every 2 days, half of the mucin agar-covered microcosms were replaced by fresh sterile ones under a flow of $N_2$ to prevent disruption of anaerobic conditions. Fourteen days after inoculation, 3 colon vessels were infected with $10^6$ WT bacteria , 3 colon vessels with $10^6$ Δ*lmo2776* and 3 were left uninfected. Lumen (10 ml) and mucin agar samples were taken at 6, 24, 48 and 72h. Mucin agar-covered microcosms were washed with sterile PBS to remove lumen bacteria. Mucin agar was removed from microcosms, homogenised and stored immediately at −20°C until further analysis.

For 16S rRNA Gene Sequencing, DNA was extracted from 1 ml of lumen samples or 250 mg of mucin agar samples using a PowerSoil DNA Isolation kit and 16S rRNA gene sequencing was analysed as decribed above. Our full 16S rRNA gene sequence data are deposited under Study ID YYYY in the QIIME-DB database (http://www.microbio.me/qiime/).

For SCFA analysis, lumen samples were diluted 1:2 in demineralized water. Acetate, propionate, butyrate, isobutyrate and isovalerate were extracted and quantified as described (*70*).

**RNA extraction and qRT**

A total of 25 ml cultures of bacteria, grown in BHI, was pelleted for 20 min at 3000g. Pellets were resuspended in 1 ml TRI Reagent (Sigma), transferred to 2-ml Lysing Matrix tubes (MP Biomedicals) and mechanically lysed by bead beating in a FastPrep apparatus (twice 45s, speed 6.5). Tubes were then centrifuged for 5 min at 8000g at 4°C to separate beads from lysates. The lysates were drawn off and transferred to a 2-ml Eppendorf tube. 200 uL chloroform was added to the lysate, shaken and incubated 10 min at room temperature, followed by centrifugation for 15 min at 13 000g at 4°C. The aqueous phase was transferred to a new tube and RNA was precipitated by the addition of 500 uL isopropanol and incubation at room temperature for 10 min. RNA was pelleted by centrifuging for 10 min at 13000 g at 4°C, washed twice with 75% ethanol. RNA pellets were resuspended in 50 to 100 uL water. For each sample, 10 ug of RNA was treated with Dnase (Turbo DNA-free, Ambion) following manufacturer’s instructions. cDNA was synthetiszed from 1 ug of RNA using QuantiTect Reverse Transcription (Qiagen) and reactions were subsenquently diluted with 180 ul of water. qRT-PCR reactions were prepared with SYBR Green master mix. Reaction cycling and quantification was carried out in an C1000 touch Thermal cycler (CFX384, Biorad). Expression levels were normalized to the *rpoB* gene. Samples were evaluated in triplicate and results represent at least three independent experiments.

**Co-cultures and culture in presence of supernatant or Lmo2776 peptide**

For co-culture assays with *B. subtilis*, a mixture of equivalent CFU ($10^7$) of *B. subtilis* and *L. monocytogenes* (WT, Δ*lmo2776* or p2776) was inoculated into 5 mL of fresh BHI and incubated at 37°C for 6 hours. Serial dilutions were plated on BHI and on Oxford media for CFU enumeration.

For culture of target in presence of *Listeria* supernatant, 25 ml of overnight culture of *Listeria* were centrifuged at 13000g and the supernatants were collected and centrifuged further to remove cells and cells debris. Supernatants were filtered through a 0.20µm pore size filter (Millipore). $10^7$ bacterial targets (*B. subtilis*, *E. coli* or *P. copri*) were inoculated at 37°C into 2.5 ml of listerial supernatant and 2.5 ml of fresh medium (BHI for *B. subtilis,* LB for *E. coli* and PYG modified for *P. copri*). At 16h after inoculation (in aerobic conditions for *B. subtilis* and *E. coli* and in anaerobic conditions for *P. copri*), cultures were serially diluted and plated on medium.

For *in vitro* assays, a peptide of 63aa (GTFWVTWGQDRHYSNYQHTKKTHRSSASNYRA TERSSWKAKNNLATAWIKSSLWGNKANWATK), corresponding to the putative mature form of Lmo2776 has been chemically synthesized (Polypeptide). $10^7$ bacteria were inoculated in absence or in presence of this peptide at 37°C into 5 ml of medium. At 16h after inoculation (in aerobic or anaerobic conditions), cultures were serially diluted and plated.

For *in vivo* assays, conventional mice were anaesthetized with an intraperitoneal injection of 75 mg ketamine $kg^{-1}$ and 5 mg xylazine $kg^{-1}$. One hundred ul of Lmo2776 peptide (1mg in 100µl distilled $H_2O$) was introduced rectally using a flexible catheter into each of 12 test mice and 100µl distilled $H_2O$ was introduced into each of 6 control mice. Feces were collected between 1 and 4 h following the introduction. Bacteria were quantified as described above.

**Immunostaining of mucins and localization of bacteria by FISH**

Mucus immunostaining was paired with fluorescent in situ hybridization (FISH), as previously described (*69, 71*). Observations were performed with a Zeiss LSM 700 confocal microscope. The software Zen 2011 version 7.1 was used to measure the distance between bacteria and epithelial cell monolayer.

#### Quantification of fecal Lcn-2 by ELISA

Fecal samples were weighted, reconstituted in PBS-0.1% Tween 20 to a final concentration of 100 mg/mL and homogenized. Samples were then centrifuged for 10 min at 14 000 g and 4°C and supernatants were collected and stored at −20°C until analysis. Lcn-2 levels were measured using DuoSet mouse Lipocalin-2/NGAL ELISA kit (R&D Systems).

#### Core genome MLST

cgMLST analysis was performed as previously described *(53).*

#### Statistical analysis

Statistical tests are reported and described in the figure legends. Differences denoted in the text as significant fall below a p-value of 0.05.

#### Figure legends

**Figure S1. Presence of *lmo2776* in *L. monocytogenes* strains and homologies with members of the Lactococcin 972 family.** (**A**) Schematic representation of the *lmo2776* genetic region. **(B)** Amino acid alignment of Lmo2776 with members of the Lactococcin 972 family: L82330 from *Lactococcus lactis* subsp. *lactis* Il1403, SP_0109 bacteriocin from *Streptococcus pneumoniae* TIGR4, SAP109 from *Staphylococcus aureus* subsp. *aureus* N315, lcn972 lactococcin 972 from *Lactococcus lactis* subsp. *lactis* and Sil from *Streptococcus iniae* SF. The conserved residues and consensus motif are indicated in color and on the bottom, respectively. **(C)** Cluster analysis based on core genome multilocus sequence typing (cgMLST) profiles of 1,096 genomes (*72*) and pattern of *lmo2774-lmo2775-lmo2776* genes presence (blue) or absence (white). The ten most frequent sublineages (SL) are highlighted. The last column corresponds to the sample source, represented by colour codes (upper left key).

**Figure S2. (A)** Relative expression of *lmo2774*, *lmo2775* and *lmo2776* in bacteria grown in stationary phase (white bars) compared to bacteria grown in exponential phase (black bars). The transcripts levels were normalised to the levels of *rpoB*, which were constant under all conditions, and then expressed relative to those of exponential phase. Results are expressed as mean ± SEM of a least 3 independent experiments and P-values were obtained using two-tailed unpaired Student's t-test (*p<0.05). **(B)** Relative expression of *lmo2774, lmo2775* and *lmo2777* in Δ*lmo2776* bacteria compared to WT bacteria, grown in exponential phase (black bars) or in stationary phase (grey bars). The transcripts levels were normalised to the levels of *rpoB*, which were constant under all conditions. Results are expressed as mean ± SEM of a least 3 independent experiments. **(C)** Growth curves of WT and Δ*lmo2776* bacteria at 37°C with shaking in BHI. **(D)** BALB/c mice were inoculated orally with $5\times10^9$ *Listeria monocytogenes* WT (EGDe) or Δ*lmo2776* bacteria. CFUs in the intestinal luminal content were assessed at 24, 48 and 72h post-infection. **(E)** BALB/c mice were inoculated intravenously with $5.10^3$ WT or Δ*lmo2776* bacteria. CFUs in the spleen and the liver were assessed at 72h post-infection. Each dot represents the value for one mouse. Statistically significant differences were evaluated by the Mann–Whitney test. (*$p< 0.05$).

**Figure S3. Effect of *Listeria* infection on mouse microbiota.** (**A**) Principal coordinates analysis of the weighted Unifrac distance matrix of conventional mice at day 0 (red) and infected with WT strain at day 1 (blue). Permanova P= 0.002. (**B**) Relative abundance of classes in conventional mice at day 0 (left) and infected with WT strain at day 1 (right). LEfSE (**C**) and histogram of the LDA scores (**D**) computed for features differentially abundant between microbiota of mice at day 0 (red) and infected with WT strain at day 1 (green). (**E**) Firmicutes/Bacteroides ratio in microbiota of mice at day 0 and infected with WT strain at day 1. Each dot represents the value for one mouse.

**Figure S4. Production of butyrate, isobutyrate, acetate and isovalerate is not modify by Lmo2776.** Levels of butyrate (**A**), isobutyrate (**B**), acetate (**C**) and isovalerate (**D**) in SHIME® vessels infected with WT (orange) or Δ*lmo2776* (red) strains or non-infected (blue) overtime. Results are expressed as mean ± SEM for 2 to 3 individual vessels.

**Figure S5.** Numbers of *B. subtilis* (**A**) and of different *Lm* strains (**B**) were quantified after 6h of co-culture. Numbers of *B. subtilis* or *E. coli* (**C**) after incubation with supernatant of WT or Δ*lmo2776* strains. Results are expressed as mean ± SEM of a least 3 independent experiments and P-values were obtained using two-tailed unpaired Student’s t-test (*$p<0.05$, ***$p<0.005$). Assessment of listerial CFUs in the liver (**D**) of germ-free (GF) C57BL/6J mice colonized or not with *P. copri*, *P. salivae* or *B. thetaiotamicron* or stably colonized with 12 bacterial species (Oligo-$MM^{12}$) for 2 weeks and then inoculated with *L. monocytogenes* WT or Δ*lmo2776* for 72h. Each dot represents one mouse.

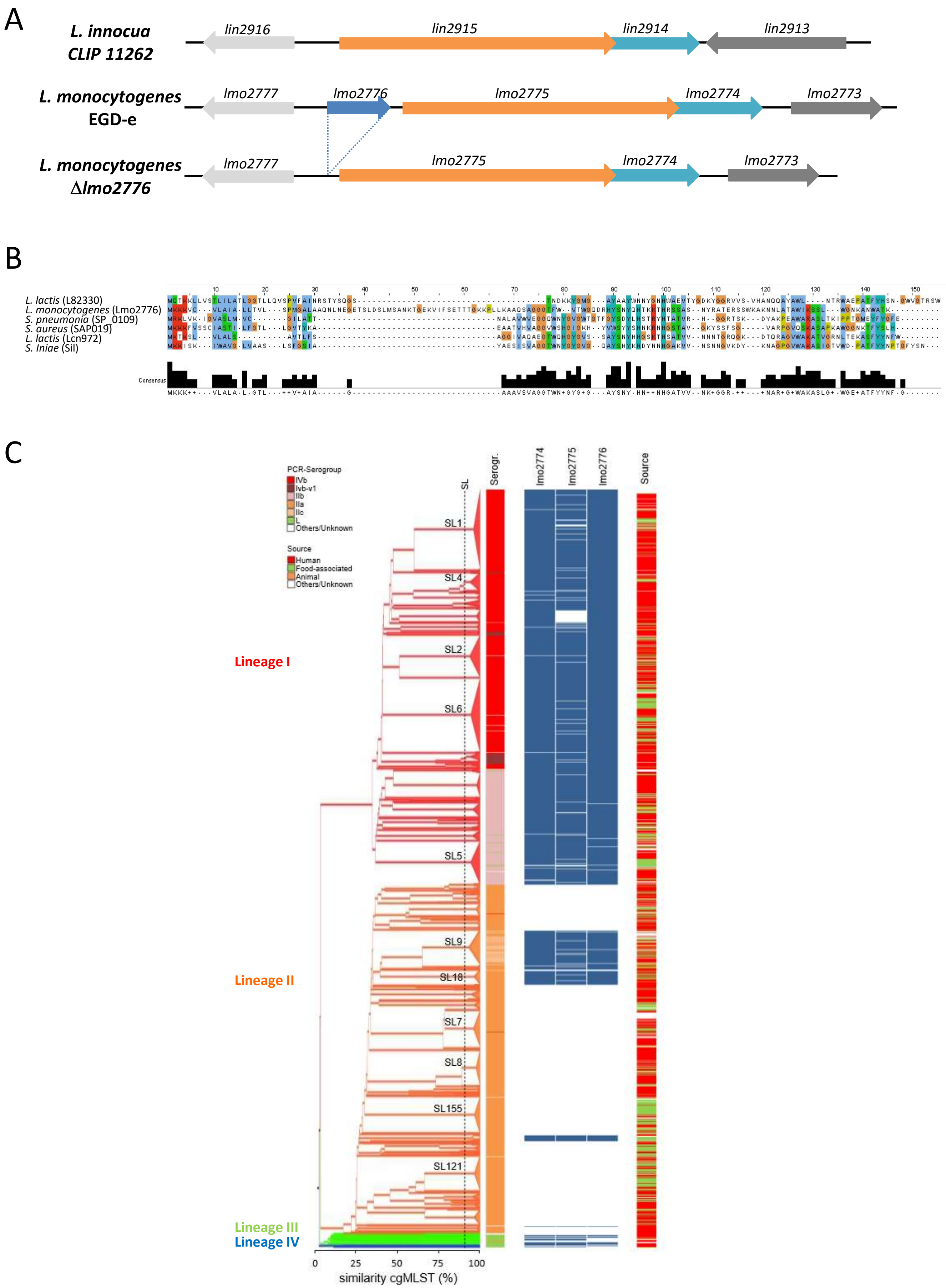
A
L. innocua CLIP 11262
lin2916
lin2915
lin2914
lin2913
L. monocytogenes EGD-e
lmo2777
lmo2776
lmo2775
lmo2774
lmo2773
L. monocytogenes Δlmo2776
lmo2777
lmo2775
lmo2774
lmo2773
B
L. lactis (L82330)
L. monocytogenes (Lmo2776)
S. pneumonia (SP_0109)
S. aureus (SAP019)
L. lactis (Lcn972)
S. iniae (Sil)
Consensus
C
PCR-Serogroup
IVb
Ivb-v1
IIb
IIa
IIc
L
Others/Unknown
Source
Human
Food-associated
Animal
Others/Unknown
SL
Serogr.
lmo2774
lmo2775
lmo2776
Source
SL1
SL4
SL2
SL6
SL5
SL9
SL18
SL7
SL8
SL155
SL121
Lineage I
Lineage II
Lineage III
Lineage IV
0
25
50
75
100
similarity cgMLST (%)

### Figure S1

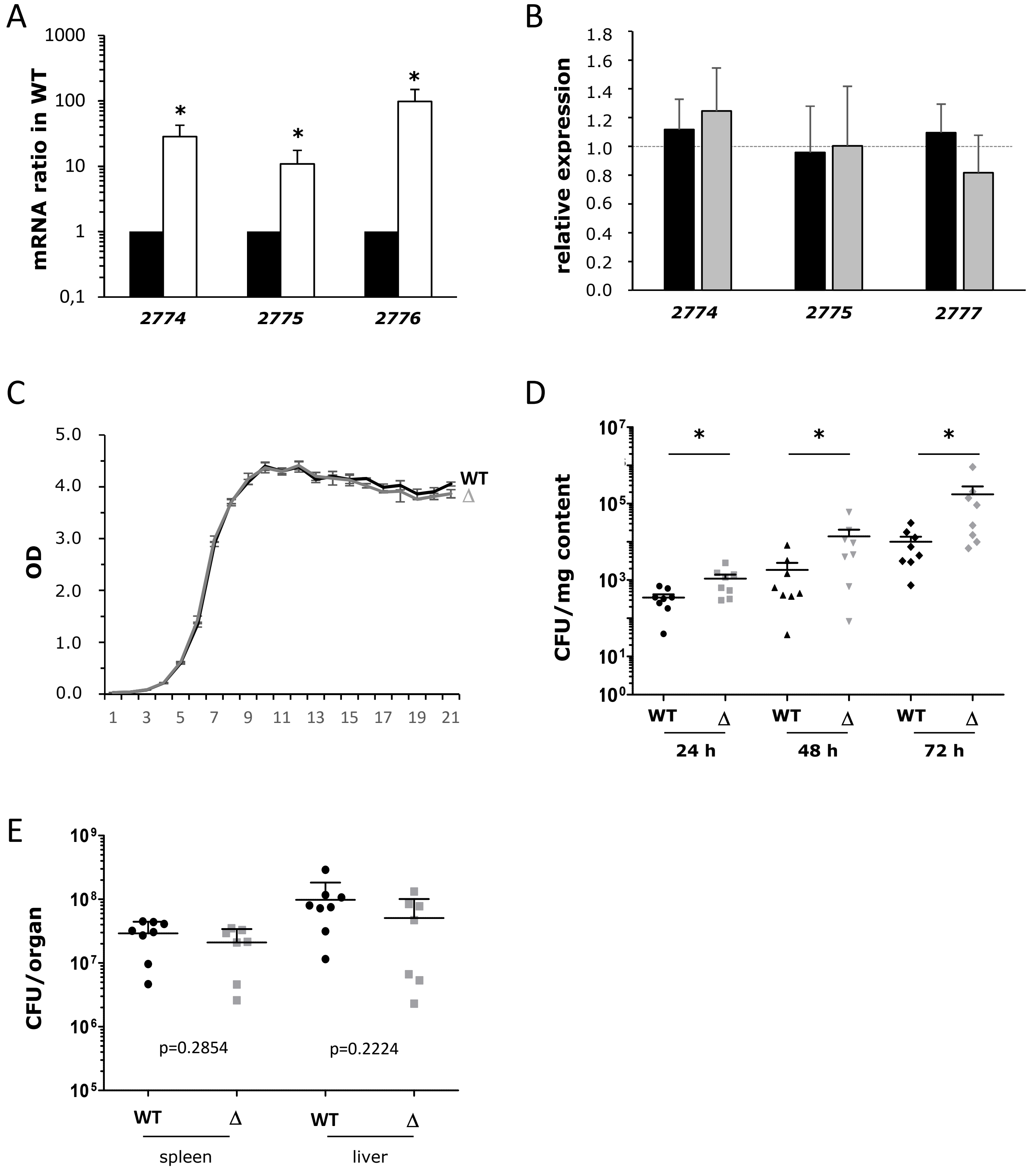

A
mRNA ratio in WT
1000
100
10
1
0,1
2774
2775
2776
B
relative expression
1.8
1.6
1.4
1.2
1.0
0.8
0.6
0.4
0.2
0.0
2774
2775
2777
C
OD
5.0
4.0
3.0
2.0
1.0
0.0
1 3 5 7 9 11 13 15 17 19 21
WT
Δ
D
CFU/mg content
WT
Δ
24 h
48 h
72 h
E
CFU/organ
p=0.2854
p=0.2224
WT
Δ
spleen
liver


**Figure S2**

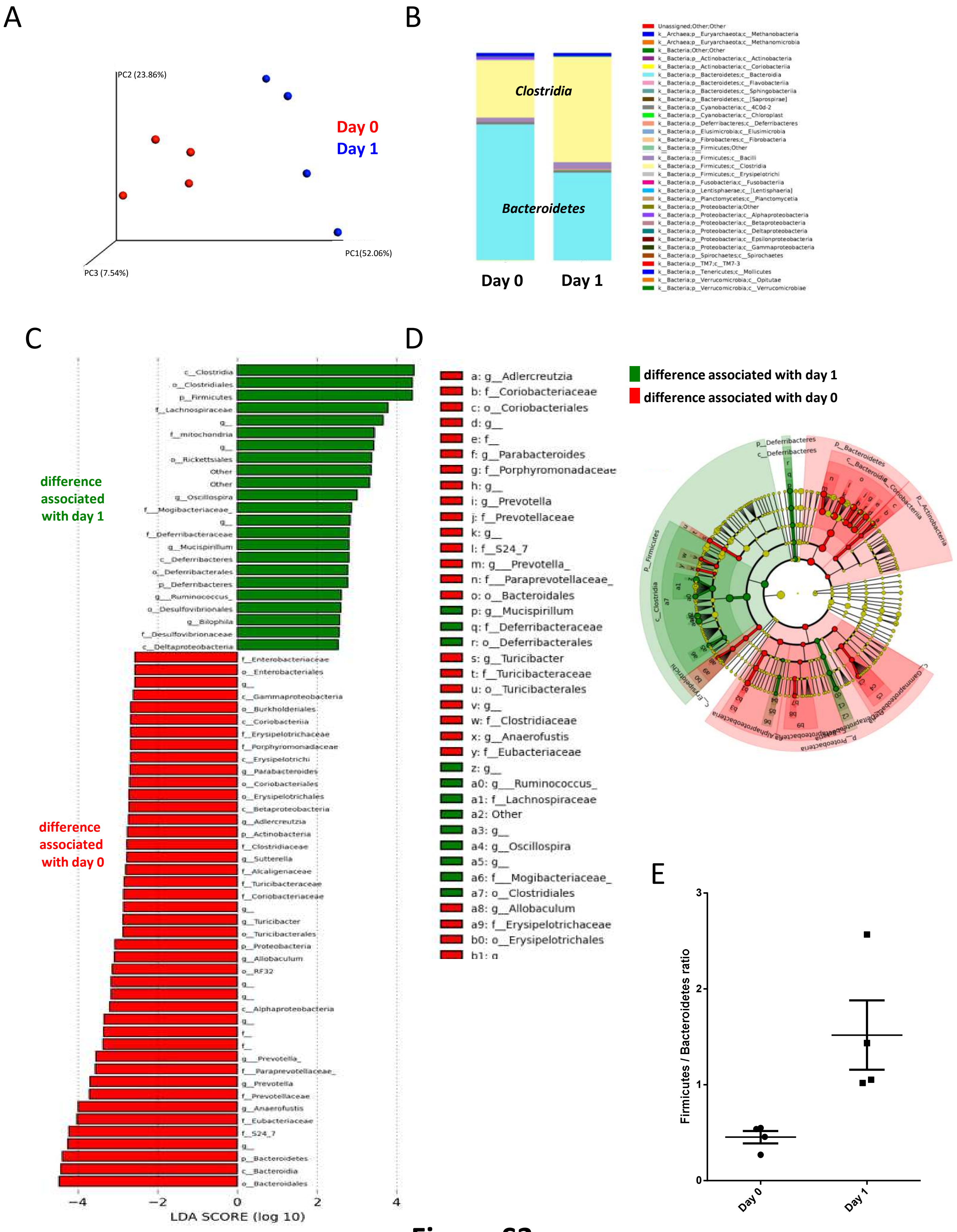
A
PC2 (23.86%)
PC1(52.06%)
PC3 (7.54%)
Day 0
Day 1
B
Clostridia
Bacteroidetes
Day 0
Day 1
Unassigned;Other;Other
k__Archaea;p__Euryarchaeota;c__Methanobacteria
k__Archaea;p__Euryarchaeota;c__Methanomicrobia
k__Bacteria;Other;Other
k__Bacteria;p__Actinobacteria;c__Actinobacteria
k__Bacteria;p__Actinobacteria;c__Coriobacteriia
k__Bacteria;p__Bacteroidetes;c__Bacteroidia
k__Bacteria;p__Bacteroidetes;c__Flavobacteriia
k__Bacteria;p__Bacteroidetes;c__Sphingobacteriia
k__Bacteria;p__Bacteroidetes;c__[Saprospirae]
k__Bacteria;p__Cyanobacteria;c__4C0d-2
k__Bacteria;p__Cyanobacteria;c__Chloroplast
k__Bacteria;p__Deferribacteres;c__Deferribacteres
k__Bacteria;p__Elusimicrobia;c__Elusimicrobia
k__Bacteria;p__Fibrobacteres;c__Fibrobacteria
k__Bacteria;p__Firmicutes;Other
k__Bacteria;p__Firmicutes;c__Bacilli
k__Bacteria;p__Firmicutes;c__Clostridia
k__Bacteria;p__Firmicutes;c__Erysipelotrichi
k__Bacteria;p__Fusobacteria;c__Fusobacteriia
k__Bacteria;p__Lentisphaerae;c__[Lentisphaeria]
k__Bacteria;p__Planctomycetes;c__Planctomycetia
k__Bacteria;p__Proteobacteria;Other
k__Bacteria;p__Proteobacteria;c__Alphaproteobacteria
k__Bacteria;p__Proteobacteria;c__Betaproteobacteria
k__Bacteria;p__Proteobacteria;c__Deltaproteobacteria
k__Bacteria;p__Proteobacteria;c__Epsilonproteobacteria
k__Bacteria;p__Proteobacteria;c__Gammaproteobacteria
k__Bacteria;p__Spirochaetes;c__Spirochaetes
k__Bacteria;p__TM7;c__TM7-3
k__Bacteria;p__Tenericutes;c__Mollicutes
k__Bacteria;p__Verrucomicrobia;c__Opitutae
k__Bacteria;p__Verrucomicrobia;c__Verrucomicrobiae
C
difference associated with day 1
difference associated with day 0
c__Clostridia
o__Clostridiales
p__Firmicutes
f__Lachnospiraceae
g__
f__mitochondria
g__
o__Rickettsiales
Other
Other
g__Oscillospira
f__Mogibacteriaceae_
g__
f__Deferribacteraceae
g__Mucispirillum
c__Deferribacteres
o__Deferribacterales
p__Deferribacteres
g__Ruminococcus_
o__Desulfovibrionales
g__Bilophila
f__Desulfovibrionaceae
c__Deltaproteobacteria
f__Enterobacteriaceae
o__Enterobacteriales
g__
c__Gammaproteobacteria
o__Burkholderiales
c__Coriobacteriia
f__Erysipelotrichaceae
f__Porphyromonadaceae
c__Erysipelotrichi
g__Parabacteroides
o__Coriobacteriales
o__Erysipelotrichales
c__Betaproteobacteria
g__Adlercreutzia
p__Actinobacteria
f__Clostridiaceae
g__Sutterella
f__Alcaligenaceae
f__Turicibacteraceae
f__Coriobacteriaceae
g__
g__Turicibacter
o__Turicibacterales
p__Proteobacteria
g__Allobaculum
o__RF32
g__
g__
c__Alphaproteobacteria
g__
f__
f__
g__Prevotella_
f__Paraprevotellaceae_
g__Prevotella
f__Prevotellaceae
g__Anaerofustis
f__Eubacteriaceae
f__S24_7
g__
p__Bacteroidetes
c__Bacteroidia
o__Bacteroidales
−4
−2
0
2
4
LDA SCORE (log 10)
D
a: g__Adlercreutzia
b: f__Coriobacteriaceae
c: o__Coriobacteriales
d: g__
e: f__
f: g__Parabacteroides
g: f__Porphyromonadaceae
h: g__
i: g__Prevotella
j: f__Prevotellaceae
k: g__
l: f__S24_7
m: g__Prevotella_
n: f__Paraprevotellaceae_
o: o__Bacteroidales
p: g__Mucispirillum
q: f__Deferribacteraceae
r: o__Deferribacterales
s: g__Turicibacter
t: f__Turicibacteraceae
u: o__Turicibacterales
v: g__
w: f__Clostridiaceae
x: g__Anaerofustis
y: f__Eubacteriaceae
z: g__
a0: g__Ruminococcus_
a1: f__Lachnospiraceae
a2: Other
a3: g__
a4: g__Oscillospira
a5: g__
a6: f__Mogibacteriaceae_
a7: o__Clostridiales
a8: g__Allobaculum
a9: f__Erysipelotrichaceae
b0: o__Erysipelotrichales
b1: g
difference associated with day 1
difference associated with day 0
p__Deferribacteres
c__Deferribacteres
p__Bacteroidetes
c__Bacteroidia
c__Coriobacteriia
p__Actinobacteria
p__Firmicutes
c__Clostridia
c__Erysipelotrichi
c__Gammaproteobacteria
p__Proteobacteria
E
Firmicutes / Bacteroidetes ratio
0
1
2
3
Day 0
Day 1

### Figure S3

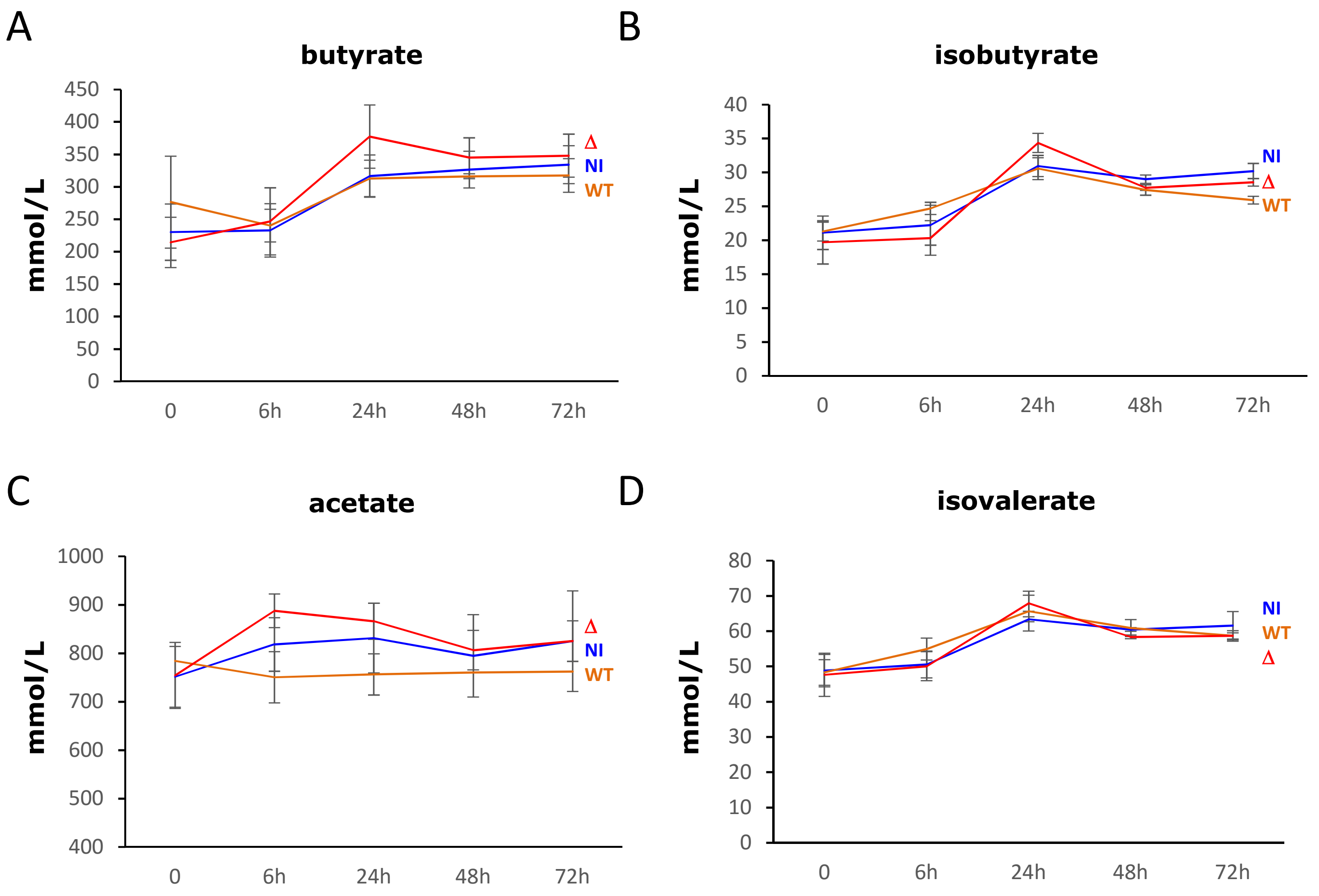
A
butyrate
mmol/L
450
400
350
300
250
200
150
100
50
0
0
6h
24h
48h
72h
Δ
NI
WT
B
isobutyrate
mmol/L
40
35
30
25
20
15
10
5
0
0
6h
24h
48h
72h
NI
Δ
WT
C
acetate
mmol/L
1000
900
800
700
600
500
400
0
6h
24h
48h
72h
Δ
NI
WT
D
isovalerate
mmol/L
80
70
60
50
40
30
20
10
0
0
6h
24h
48h
72h
NI
WT
Δ

**Figure S4**

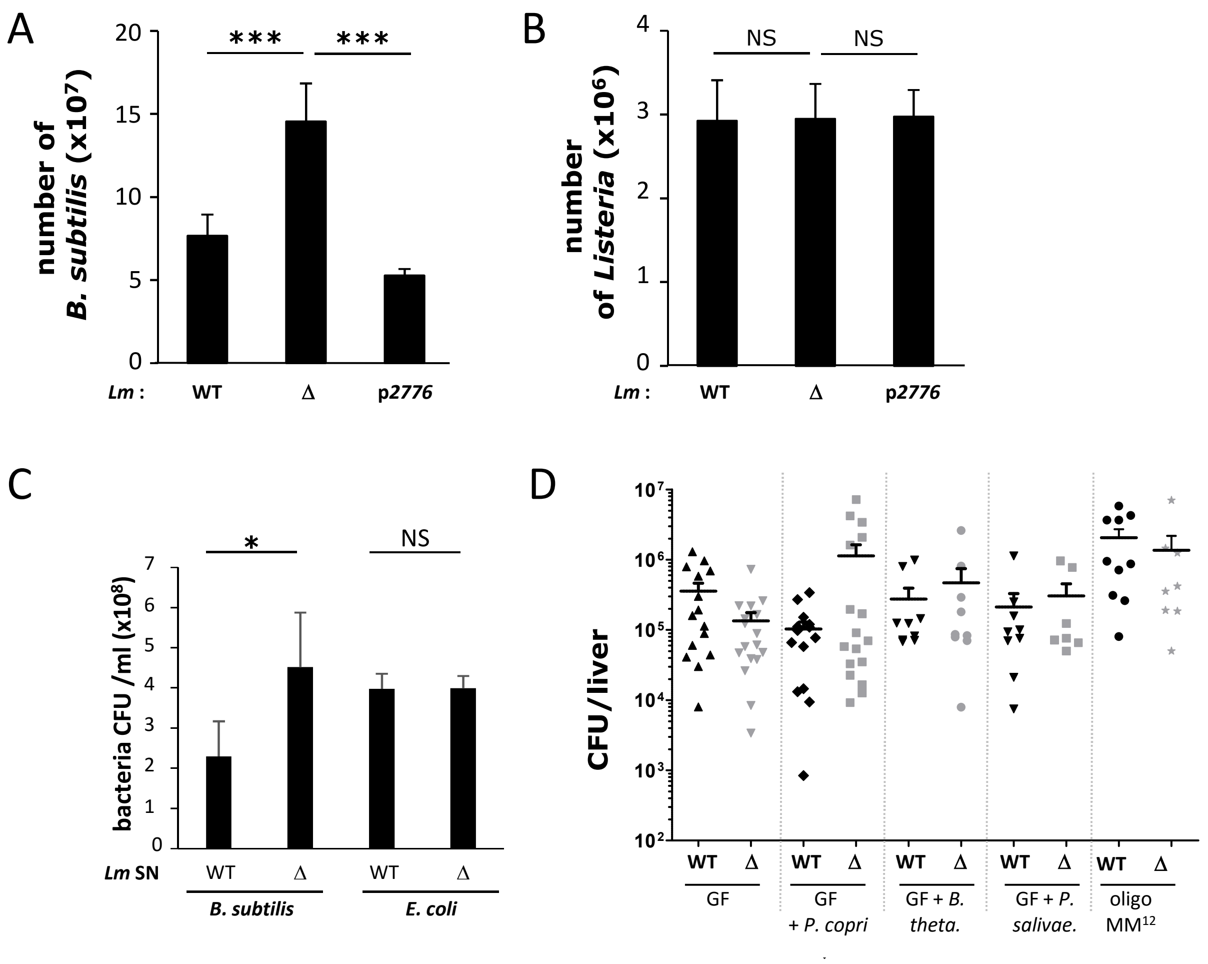

A
B
C
D
number of B. subtilis (x10^7)
number of Listeria (x10^6)
***
NS
Lm :
WT
Δ
p2776
bacteria CFU /ml (x10^8)
*
Lm SN
B. subtilis
E. coli
CFU/liver
GF
GF + P. copri
GF + B. theta.
GF + P. salivae.
oligo MM12


### Figure S5

#### Table S1 : Strains and plasmids used in this study

| Strains | Characteristics | Collection no. | Source or reference |
|---|---|---|---|
| EGD-e | *Listeria monocytogenes* WT strain | BUG 1600 | Mackanes et al 1964 |
| EGD-e Δ*lmo2776* | *Listeria monocytogenes* EGD-e *lmo2776* deletion mutant | BUG 3713 | this study |
| EGD-e Δ*lmo2776* p*lmo2776* | pAD-*lmo2776* chromosomally integrated in Δ*lmo2776* | BUG 3717 | this study |
| F2365 | *Listeria monocytogenes* strain associated with the 1985 listeriosis outbreak in California | BUG 3012 | |
| 10403S | *Listeria monocytogenes* WT strain | BUG 1361 | |
| *L. innocua* | *L. innocua* Clip11262 | BUG 499 | |
| *Bacillus subtilis* | *Bacillus subtilis* 168trpC2 | BUG 748 | |
| *Escherichia coli* Top10 | F- mcrA Δ(mrr-hsdRMS-mcrBC) ϕ80*lacZ* ΔM15Δ*lac* X74 *rec* A1 araD139 Δ(ara-leu) 7697 *galU galK rpsL* (StrR) endA1 nupG λ- | | Invitrogen |
| *Bacteroides thethaiotamicron* | strain from the Institut Pasteur Collection | CIP 104206T | |
| *Lactococcus lactis* | *Lactococcus lactis* IL14103 | BUG 1801 | |
| *Lactococcus lactis* | *Lactococcus lactis* M1363 | BUG 3029 | |
| *Enterococcus faecalis* | *Enterococcus faecalis* | BUG 3402 | ATCC 700802 |
| *Akkermensia muciniphila* | ATCC BA-835 | | ATCC |
| *Prevotella copri* | DSMZ 18205 | BUG 4034 | DSMZ |
| *Prevotella salivae* | DSMZ 15606 | BUG 4184 | DSMZ |
| *Prevotella oris* | strain from the Institut Pasteur Collection | CIP 104480T | |
| *Prevotella nigrescens* | strain from the Institut Pasteur Collection | CIP 105552T | |
| *Prevotella pallens* | strain from the Institut Pasteur Collection | CIP 105551T | |
| *Prevotella corporis* | strain from the Institut Pasteur Collection | CIP 105107T | |
| *Prevotella melanogenica* | strain from the Institut Pasteur Collection | CIP 104591 | |
| *Prevotella bivia* | strain from the Institut Pasteur Collection | CIP 105105T | |

| Plasmids | Characteristics | Collection no. | Source or reference |
|---|---|---|---|
| pMAD | shuttle vector used for creating plasmid for mutagenesis | | *59* |
| pAD | site-specific integration vector used for complementation | | *61* |
| p*lmo2776* | *lmo2776* complementation plasmid | BUG 4020 | this study |

#### Table S2 : Oligonucleotide primers used in this study

| **Primer** | **Sequence** |
|---|---|
| Oligonucleotides used to create deletion mutant | |
| Lmo2776-DelA | GGAAGATCTACATCCTTCACAGGGAAATG |
| Lmo2776-DelB | TGTATTCTCCTCTCTTTCAAATTAA |
| Lmo2776-DelC | TTAATTTGAAAGAGAGGAGAATACATTCCTATAAAGCTAAGAAATATTTC |
| Lmo2776-DelD | CGGCCATGGAGCAAAGTCATAAGTAACGGGATAT |
| Sequencing insert in pAD-based plasmid | |
| pPL2-Fw | TTCGACCCGGTCGTCGGTTC |
| pPL2-Rv | CTTAGACGTCATTAACCCTCAC |
| Verification of pAD integration in the *Listeria* chromosome | |
| NC16 | GTCAAAACATACGCTCTTATC |
| PL95 | ACATAATCAGTCCAAAGTAGATGC |
| Creation of the p*lmo2776* : lux plasmid | |
| Lmo2776 P1 | CCATCTCGAGAAAAAACTCTCCTGAATAATTTATTTATC |
| Lmo2776 P2 | CATTGTATTCTCCTCTCTTTCAAATTAA |
| qRT-PCR | |
| lmo2774 1 | GTGTTAGAAAACTTATCCATTACAGG |
| lmo2774 2 | ATCATCCAAGTTGCCTGTTGGTTCG |
| lmo2775 1 | GGTCTTTAAAGGTTTATGACTTTGG |
| lmo2775 2 | CCCATATACGATTAACTAATAACCG |
| lmo2776 RT1 | GTATGTGTTTTAGCGATAGCA |
| lmo2776 RT2 | ATAATGTCTATCTTGTCCCC |
| lmo2777 RT1 | CTACCACGGAAATGATCGCC |
| lmo2777 RT2 | GCACACTAAAGGAGGTAGCG |
| rpoB R | ATGTTTGGCAGTTACAGCAGCACC |
| rpoB F | GCGAACATGCAACGTCAAGCAGTA |
| q-PCR | |
| *Prevotella* 16S F | CACRGTAAACGATGGATGCC |
| *Prevotella* 16S R | GGTCGGGTTGCAGACC |
| Universal 16S F | ACTCCTACGGGAGGCAGCAGT |
| Universal 16S R | ATTACCGCGGCTGCTGGC |

| | |
|---|---|
| Am_fwd | CAGCACGTGAAGGTGGGGAC |
| Am_rev | CCTTGCGGTTGGCTTCAGAT |
